# The role of serotonin in the orbitofrontal cortex in alcohol consumption

**DOI:** 10.64898/2026.08.19.745752

**Authors:** Jessica A. Wojick, Sofia Neira, Kristen Boyt, Christina Stanhope, Sarah Y. Wu, Alexandra M. Weir, Meghan Flanigan, Verginia C. Cuzon Carlson, Jobe L Ritchie, Kathleen A. Grant, Thomas L. Kash, Melanie M. Pina

**Author notes:** co-corresponding authors, CORRESPONDING AUTHORS’ EMAIL ADDRESSES, and.

## Abstract

Binge alcohol drinking is a public health concern that can dramatically increase the risk for development of alcohol use disorder (AUD). Continued alcohol drinking in the face of negative consequences is another key feature of AUD. A better understanding of the neural circuitry that regulates these behaviors could provide insight as to novel treatments for AUD. Serotonin is a neurotransmitter that has been implicated in alcohol consumption in both human studies and animal models. The orbitofrontal cortex (OFC) is a brain region that both receives serotonergic input from the dorsal raphe and has been implicated in AUD. However, how volitional alcohol consumption impacts serotonin signaling within the OFC and how this contributes to alcohol related behaviors is unknown. Here, we show that a history of alcohol consumption alters the ability of 5-HT to hyperpolarize OFC pyramidal neurons in mice and monkeys. Consistent with this, a history of binge alcohol consumption decreases the expression of the 5-HT_1A_ but not 5-HT_2A_ receptor in the OFC from mice. Next, we show that deletion of the 5-HT_1A_ receptor from the OFC increased alcohol intake and preference in male, but not female mice. Finally, we found that 5-HT_1A_ receptor deletion led to increased quinine-adulterated alcohol intake, a measure of aversion-resistant drinking, in both male and female mice. Altogether, we identified serotonin signaling in the OFC as key target for modulation of binge and compulsive alcohol consumption.

**SIGNIFICANCE STATEMENT:** Binge alcohol drinking is a public health concern that increases the risk for development of alcohol use disorder (AUD). To develop novel treatments for AUD, there is a need to better understand the neural circuitry that regulates binge alcohol consumption. Serotonin (5-HT) is a neuromodulator implicated in alcohol consumption in human studies and animal models. The orbitofrontal cortex (OFC) is a brain region that both receives 5-HT input and has been implicated in AUD. Here, we show that a history of alcohol consumption alters 5-HT signaling in the OFC of mice and monkeys and that alterations in 5-HT signaling increases alcohol consumption. Altogether, we identified 5-HT signaling in the OFC as a key target for modulating alcohol consumption.

## INTRODUCTION

A rapid increase in the incidence of binge drinking constitutes a leading cause of mortality and economic burden globally (Dwyer-Lindgren et al., 2016, 2016; Esser, 2014; Naimi et al., 2003; Sacks et al., 2015; Stahre, 2014; Weerakoon et al., 2021). Binge drinking is defined as the rapid consumption of alcohol leading to a blood alcohol concentration of 0.08 g/dL or greater over two hours (Courtney and Polich, 2009; Substance Abuse and Mental Health Services Administration (US) and Office of the Surgeon General (US), 2016). Animal models of binge drinking allow for the probing of neural adaptations and molecular mechanisms that are impacted by alcohol and their causal role in regulation of behavior (Bedendo et al., 2017; Crabbe et al., 2011; Gowin et al., 2017; Hingson et al., 2017; Naimi et al., 2003; Sprow and Thiele, 2012). These adaptations may contribute to alcohol misuse and the development of alcohol use disorder (AUD). Mouse models of binge drinking, including the Drinking in the Dark (DiD) model, result in high levels of alcohol intake (≥80mg/dl) that provides the opportunity to identify the cellular and molecular changes driven by alcohol exposure (Crabbe et al., 2011; Rhodes et al., 2005; Thiele et al., 2014). The inability to control alcohol consumption despite adverse consequences is an aspect of AUD that may be driven by distinct signaling to alcohol consumption alone and has been modeled in rodents via aversion resistant drinking of alcohol altered with quinine, a bitter substance. Studying both binge-like alcohol consumption and aversion-resistant alcohol drinking provides an opportunity to understand converging mechanisms that regulate both.

Rodent models allow for precise genetic and neural manipulations to probe brain function, however, the similarities to the human brain are mainly conserved to the more primitive regions, with the prefrontal cortex being the most different. Non-human primates provide unique benefits in drug abuse research, as their cortex and decision making more closely models humans. An additional benefit is that non-human primate models are often years-long voluntary access models that allows for the separation of low, moderate and high drinkers, which can also model AUD. Variability in alcohol consumption patterns in primates may more closely represent the cellular changes present in humans with AUD (Baker et al., 2017, 2014; Grant et al., 2008). While there are some pharmacological treatment options available for AUD, their use is limited, potentially due to side effects (Akbar et al., 2018). Therefore, there is a critical need to better understand the neural mechanisms that drive binge drinking and AUD to better target and treat the neuropathology underlying this disorder.

Studies in rodents and non-human primate have confirmed the relevance and importance of serotonin 5 hydroxy-tryptamine (5-HT) signaling in alcohol intake and withdrawal (Beck et al., 1983; Hillmer et al., 2014; Pivac et al., 2004; Underwood et al., 2007; Wood et al., 2022). 5-HT is a modulatory neurotransmitter that plays a role in regulating a range of behaviors. 5-HT1A and 2A receptors (5-HT_1A_R; 5-HT_2A_R) are common and functionally opposing receptors for 5-HT modulation. 5-HT_1A_R is inhibitory (Gi/o coupled) while 5-HT_2A_R is Gq-coupled and excitatory. Interestingly, acute alcohol exposure in humans drives an increase in 5HT metabolites in the urine, suggesting that there is increased 5-HT in the brain. This is supported by studies in animal models in which acute alcohol can increase 5-HT in discrete brain regions. These alcohol driven increases are reduced over time, suggesting a form of molecular tolerance (Lovinger, 1997). Polymorphisms in human genes involved in 5-HT production and transport have been associated with risk for AUD (Marcinkiewcz, 2015). The dorsal raphe is the major source of 5-HT in the brain and sends serotonergic projections to many downstream brain regions, including to the orbitofrontal cortex (OFC), a projection that has been implicated in reward encoding (Miyazaki et al., 2020; Roberts, 2011; Zhou et al., 2015). The OFC is a brain region in the prefrontal cortex that encodes motivational value to regulate outcome-based decision making and to control goal-directed behavior (Gourley et al., 2013; Gremel and Costa, 2013).

Dysfunction of the OFC and 5-HT neuromodulation has been implicated in AUD and other substance use disorders (Dom et al., 2005; Everitt et al., 2007; Moorman, 2018; Schoenbaum and Shaham, 2008; Volkow and Fowler, 2000). Further, alcohol exposure modulates 5-HT in the OFC (Nimitvilai et al., 2018, 2017, 2016), and inhibitory neurons in the OFC can suppress binge drinking (Gimenez-Gomez et al., 2025), but it is unknown how binge-like alcohol drinking impacts 5-HT signaling in the excitatory pyramidal neurons of the OFC nor how 5-HT impacts alcohol intake and preference. Here, we show that 5-HT signaling is altered in the OFC of mouse and non-human primates following high levels of alcohol consumption. In mice, binge-like alcohol consumption downregulates the 5-HT_1A_ receptor, but this correlates to alcohol in sex-specific ways. Furthermore, the 5-HT_1A_ receptor is necessary for binge-like alcohol consumption, including aversion-resistant drinking, but does not modulate consumption of non-alcohol rewarding or aversive substances. Therefore, modulating serotonin signaling, specifically by enhancing 5-HT_1A_ receptor function may represent an exciting new direction for development of treatments for AUD.

## METHODS & MATERIALS

### Mice

Mice were maintained on a 12-hour reverse light/dark cycle in temperature- and humidity-controlled facilities and given *ad libitum* access to food (Prolab IsoPro RMH 3000, LabDiet) and water. Male and female C57BL/6J mice (JAX stock #000664) were obtained from Jackson Laboratory at 8 weeks of age for slice electrophysiology and fluorescent in-situ hybridization experiments. Experimental mice were singly housed at least three days before initiation of the DiD procedure and remained singly housed throughout the completion of the experiment. *Htr1a^fl/^*^fl^ mice (JAX stock #029608) were backcrossed for several generations to C57BL/6J mice before use in genetic knockdown experiments and obtained from our in-house breeding colony. Experimental mice were group-housed until six weeks following surgery and were subsequently single-housed to allow for accurate monitoring of fluid consumption for the duration of the experiment. All procedures followed the National Institutes of Health Guide for the Care and Use of Laboratory Animals and were approved by the University of North Carolina-Chapel Hill School of Medicine Institutional Animal Care and Use Committee.

### Monkeys

Fifteen young adult rhesus macaques (*Macaca mulatta*) born and raised at the Oregon National Primate Research Center were used in this study. The monkeys were from 2 cohorts (cohorts 16 and 17 in the Monkey Alcohol Tissue Research Resource^1^ (MATRR.com). Cohort 16 provided tissue from 2 females and 4 males (6-7 years of age at necropsy) and cohort 17 provided tissue from 6 females and 5 males (7-8 years of age at necropsy). The monkeys were housed in individual cages that provided horizontal pairing with a social partner for 1-2 hrs/day. All monkeys completed an alcohol self-administration protocol (Baker et al, 2014) that differed in the amount of time they remained in the protocol. Briefly, monkeys were first trained to operate a panel to self-administer water and then were gradually induced to self-administer increasing concentrations of ethanol (0.5, 1.0, and 1.5 g/kg ethanol, 4% w/v) over three consecutive 30-day sessions, as previously described (Grant et al., 2008; Helms et al., 2014). After training, monkeys were given “open access” to choose between drinking water and 4% ethanol (w/v) in water for 22 hours a day, 7 days a week. Cohort 16 remained in open access for 216 days (approximately 7 months) and cohort 17 for 425 days (approximately 14 months). All procedures were conducted in accordance with the Guide for the Care and Use of Laboratory Animals and approved by the Oregon National Primate Research Center IACUC. Average daily ethanol intakes (g/kg) in the open access phase determined each monkeys placement into one of four drinking categories (low, binge, heavy and very heavy) based on mathematical modeling (Baker et al., 2023, 2014). As noted in previous publications: Very Heavy Drinking animals are defined by having an average daily ethanol intake exceeding 3 g/kg and have > 10% of open access days with an ethanol intake exceeding 4 g/kg; Heavy Drinkers display a daily ethanol consumption of > 3 g/kg for > 20% of days; Binge Drinkers have an ethanol consumption of > 2 g/kg for > 55% of days and a recorded blood ethanol concentration (BEC) > 80 mg/dl at least once per year. All other remaining animals are defined as Low Drinkers. These groups are stable over time (Baker et al. 2023). For the purposes of data analysis, heavy drinkers and very heavy drinkers were combined into one “heavy drinking” (HD) category for 3 total categories: Low drinkers (LD), Binge drinkers (BD) and HD.

### Stereotaxic Surgery

Adult *Htr1a^fl/^*^fl^ mice (>7 weeks of age) were anesthetized with isoflurane (1–3%) in oxygen (1–2 L/min) and positioned in a stereotaxic frame using ear cup bars (Kopf Instruments). The scalp was sterilized with 70% ethanol and betadine and a vertical incision was made before using a drill to burr small holes in the skull directly above the injection targets. Using a 1 µl Neuros Hamilton Syringe (Hamilton, Inc.), a control GFP virus AAV5-CAMKII-GFP (UNC AV4621B; titer: 4 x 10^12^ vg/mL) or a Cre-encoding virus AAV5-CAMKII-Cre-GFP (UNC AV6450; titer: 5 x 10^12^ vg/mL) was then microinjected into the OFC (mm relative to bregma: AP: 2.5, ML: ± 1.5, DV: −2.5) at a total volume of 300 nL per hemisphere. Mice were left to recover and for viral expression at least six weeks before beginning behavioral experiments.

### Drinking in the dark

Mice were given free access to both water and 20% (w/v) ethanol bottles in the home cage from 10:00 A.M. to 12:00 P.M. on Mondays, Tuesdays, and Wednesdays, and from 10:00 A.M. to 2:00 P.M. on Thursdays. At all other times, mice were given access to water alone. Water and ethanol bottles were weighed at the 2 h time point on Mondays, Tuesdays, and Wednesdays, and at the 2 h and 4 h time point on Thursdays. Ethanol and water bottle positions were alternated daily to account for any inherent side preference in the animals. This weekly DiD access schedule was repeated for 3 weeks total, after which mice remained abstinent until behavioral testing. Confirmation of binge levels of intoxication was performed by measuring the blood ethanol concentration of tail blood collected at the end of a 4 h DiD drinking session using the AM1 Analox Analyzer (Analox Instruments).

### Drinking behavioral experiments

To examine the impact of knockdown of 5-HT_1A_ receptor in the OFC on consumption of appetitive and aversive substances, mice were tested for drinking and preference of sucrose, quinine, and quinine-adulterated ethanol. The same mice that underwent three weeks of DiD and were tested for ethanol preference were given access to 2% sucrose in water for four hours. Bottles were weighed for intake measures two and four hours after being placed. A separate cohort of mice were tested for quinine (100 μM) and quinine-adulterated (100 μM) ethanol (20%) intake and preference, with measurements of bottle weight taken two and four hours after being placed.

### Fluorescent in situ hybridization

To prepare tissue for *in situ* hybridization, mice were anesthetized with isoflurane and rapidly decapitated. Brains were extracted and flash frozen on dry ice before being stored at −80 °C until sectioning. 18 µm coronal sections were collected with a Leica CM3050 S cryostat (Leica Microsystems) and mounted directly on slides. ISH was performed using the RNAscope Fluorescence Multiplex Assay Kit (Advanced Cell Diagnostics) according to the manufacturer’s instructions to fluorescently label mRNA for mouse serotonin receptor 2a (MmHtr2a-C3, cat. #401291-C3) and mouse serotonin receptor 1a (MmHtr1a-C2, Cat no. 312301-C2). Following ISH, the slides were cover slipped with Prolong-Diamond MJHMounting Medium with DAPI (Thermo Fisher Scientific). for analysis, a minimum of 5 puncta per cell was used as the criteria for a positive cell for any one mRNA marker. 1–4 images were analyzed per animal per experiment.

### Mouse slice electrophysiology

Whole-cell patch-clamp recordings were obtained from the lateral orbitofrontal cortex (within 2.3-2.8 AP in stereotaxic coordinates from Bregma). Mice were sacrificed 24-hrs or 7-10 days following DID or after water only exposure. Following rapid decapitation, brains were extracted and immersed in a chilled sucrose cutting solution (in mM: 194 sucrose, 20 NaCl, 4.4 KCl, 2 CaCl_2_, 1 MgCl_2_, 1.2 NaH_2_PO_4_, 10 D-glucose, and 26 NaHCO) oxygenated with 95% O_2_ /5% CO_2_. Coronal slices (250 µM) containing the OFC were prepared on a vibratome (VT1200S, Leica) and transferred to oxygenated aCSF (in mM: 124 NaCl, 4.4 KCl, 1 NaH_2_PO_4_, 1.2 MgSO_4_, 10 D-glucose, 2 CaCl_2_, and 26 NaHCO_3_) and maintained at 35°C for a 30 min recovery period prior to recording. A potassium gluconate-based internal solution (in mM: 135 K-gluconate, 5 NaCl, 2 MgCl_2_, 10 HEPES, 0.6 EGTA, 4 Na_2_-ATP, 0.4 Na_2_-GTP) was used for intrinsic excitability experiments and during recordings of resting membrane potential (RMP) during tests of 5-HT bath application. Recording pipettes (2–4 MΩ) were pulled from thin-walled borosilicate glass capillaries using a P95 pipette puller (Sutter Instrument). Signals were acquired using an Axon MultiClamp 700B amplifier (Molecular Devices), digitized at 10 kHz, filtered at 3 kHz, and analyzed in pClamp 10.7 (RMP). Series resistance (R_s_) was monitored without compensation, and changes in R_a_ exceeding 25% were used as exclusion criteria. 24 cells (n = 7 H2O, n = 11 24h, n = 7 7-10d) from male mice were recorded (n = 4 H2O males, 6 24h males, 3 7-10d males). 28 cells (n = 9 H2O, n = 12 24h, n = 7 7-10d) from female mice were recorded (n = 5 H2O females, 6 24h females, 4 7-10d females).

### Monkey slice electrophysiology

The necropsy procedure was performed as previously described (Davenport et al., 2014, Pleil et al., 2015). Brains were rapidly removed and a block of tissue containing the OFC was isolated. The tissue block was placed in a conical tube of ice-cold oxygenated aCSF (in mM: 124 NaCl, 4.5 KCl, 1 MgCl_2_, 26 NAHCO_3_, 1.2 NAH_2_PO_4_, 10glucose and 2 CaCl_2_) continuously aerated with 95% O_2_ /5% CO_2_ and transported on ice for slicing. The tissue was then transferred to ice-cold cutting solution containing (in mM) 194sucrose, 30 NACl, 4.5 KCl, 1 MgCl_2_, 26 NAHCO_3_, 1.2 NaH_2_PO_4_ and 10 glucose, aerated with 95% O_2_ /5% CO_2_. 300 µM thick coronal slices were obtained with a ceramic blade (Campden Instruments Ltd., Lafayette, IN, USA) attached to a VT 1200S vibratome tissue slicer (Leica Biosystems, Wetzlar, Germany). Slices were equilibrated in aCSF at 33°C for 1 hour and then transferred to room temperature until experimental use. Slices were transferred to a recording chamber fixed to the stage of an upright microscope, stabilized by an overlying platinum ring and continuously perfused with solution maintained at 28-32°C. Individual OFC pyramidal neurons were identified using a 40X water immersion objective and images were displayed on a computer monitor, aiding the navigation and placement of the recording pipette. Patch pipettes were pulled from borosilicate glass capillaries (1.5 mm outer diameter, 0.86 mm inner diameter, Sutter Instruments Co., Novato, CA, USA) and filled with a potassium gluconate based intracellular solution as used above. Recordings were made using a MultiClamp 700B amplifier (Molecular Devises, Foster City, CA, USA). Whole-cell membrane currents were digitized at 20 kHz, filtered at 2 kHz, and analyzed with Clampit 11.4.2 software (Molecular Devices, Sunnyvale, CA, USA). 15 cells (n = 7 LD, n = 6 HD, n = 2 BD) from male monkeys were recorded (n = 2 LD males, (2 HD and 1 VHD male) combined to 3 HD males, 1 BD males). 8 cells (n = 3 LD, n = 1 HD, n = 4 BD) from female monkeys were recorded (n = 3 LD females, 1 HD females, 3 BD females).

### Experimental design and statistical analyses

Statistical analyses were performed using GraphPad Prism 10 software. One-way ANOVAs with a Bonferroni correction for multiple comparisons were used to analyze electrophysiology data from mice. Two-way ANOVAs with a Sidak’s correction for multiple comparisons were used to analyze electrophysiology data from monkeys and behavioral data from mice. The lifetime ethanol consumption for each monkey was plotted vs. the 5-HT-induced effect on membrane potential to determine if there was a relationship between the total consumption of alcohol and modulation of OFC neurons. Significance was set at p < 0.05. In all figures, significance corresponds to the following: * p < 0.05.

## RESULTS

### Repeated ethanol drinking reduces 5-HT-induced inhibition of the OFC in male mice

To examine the impact of repeated cycles of ethanol drinking on serotonin signaling, we gave mice access to 20% w/v ethanol using the two-bottle drinking in the dark (DiD) method over three weeks. We performed whole-cell current-clamp electrophysiological recordings from pyramidal neurons in the OFC from male and female mice that were ethanol naïve (water), 24 hours after the last ethanol binge session (24h) or 7-10 days abstinent from ethanol (7-10d) and recorded membrane potential in current clamp before and after bath application of 5-HT as previously described (Flanigan et al., 2023a; Pati et al., 2023; Taxier et al., 2025). There was no effect on Resistance (Table 1) or capacitance (Table 2). Neurons from water control males were hyperpolarized in the presence of 5-HT, consistent with previous reports. This 5-HT-induced hyperpolarization was not present in neurons from 24h and 7-10d male mice, which displayed significantly less change from baseline membrane potential (Fig. 1A-C). In contrast to the male mice, we did not find a consistent effect of 5-HT on membrane potential in female mice (Fig. 1D, E).

**Figure 1.**
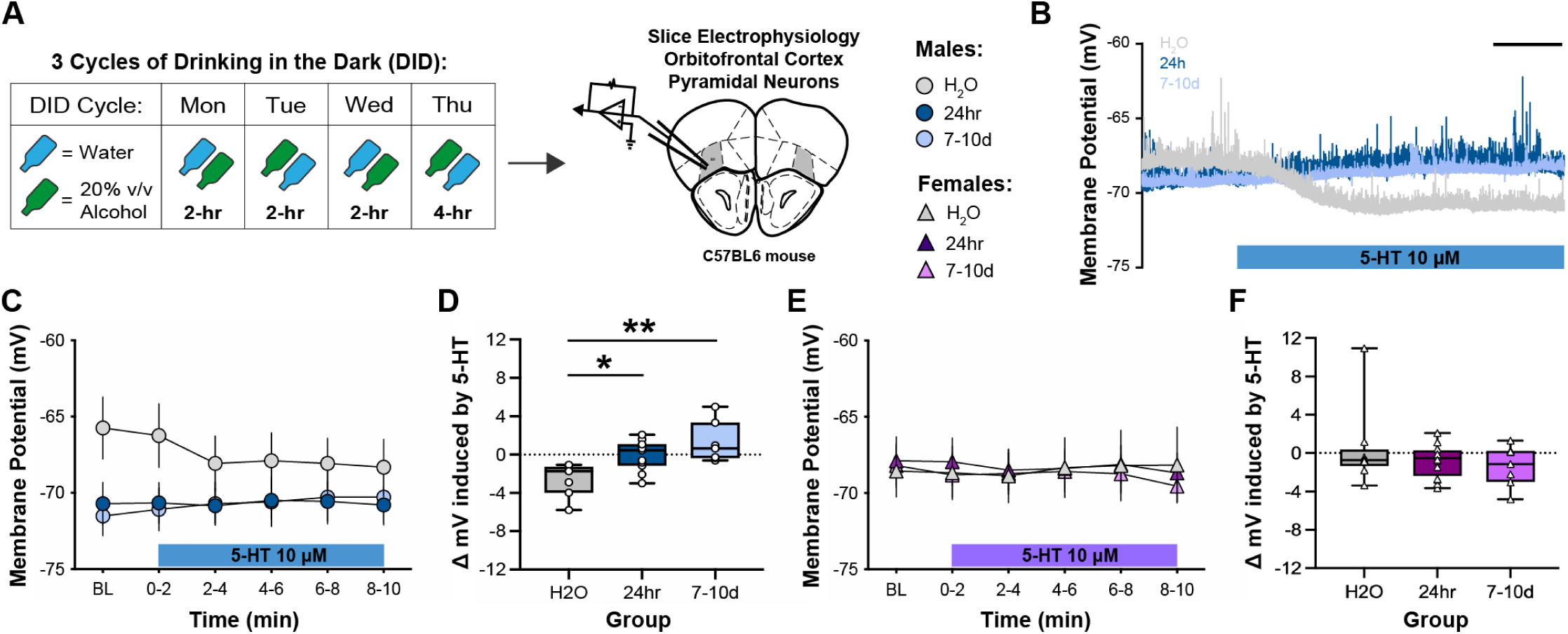
Binge-like ethanol consumption prevents 5-HT induced hyperpolarization of the OFC in male mice. **A.** Individual example electrophysiological traces from male mice exposed to water drinking or ethanol drinking 24 hours or 7-10 days after last binge session showing membrane potential in mV 1.5 min before and 4.5 min after application of serotonin (5-HT). Scale: 1 min. **B.** Average membrane potential of H2O, 24h, and 7-10d post binge-like ethanol drinking male mice at baseline and 10 minutes post-application of 5-HT. **C.** Change in membrane potential induced by 5-HT application in male mice (One-way ANOVA, F_(2,22)_ = 8.28, p = 0.002; Bonferroni-corrected pairwise comparisons, H2O vs. 24h: p = 0.026, H2O vs. 7-10d: p = 0.002, 24h vs. 7-10d: p = 0.419). **D.** Average membrane potential of H2O, 24h, and 7-10d post binge-like ethanol drinking female mice at baseline and 10 minutes post-application of 5-HT. **E.** Change in membrane potential induced by 5-HT application in female mice (One-way ANOVA, F_(2,25)_ < 1). 24 cells (n = 7 H2O, n = 11 24h, n = 7 7-10d) from male mice were recorded (n = 4 H2O males, 6 24h males, 3 7-10d males). 28 cells (n = 9 H2O, n = 12 24h, n = 7 7-10d) from female mice were recorded (n = 5 H2O females, 6 24h females, 4 7-10d females).

**Table 1.**
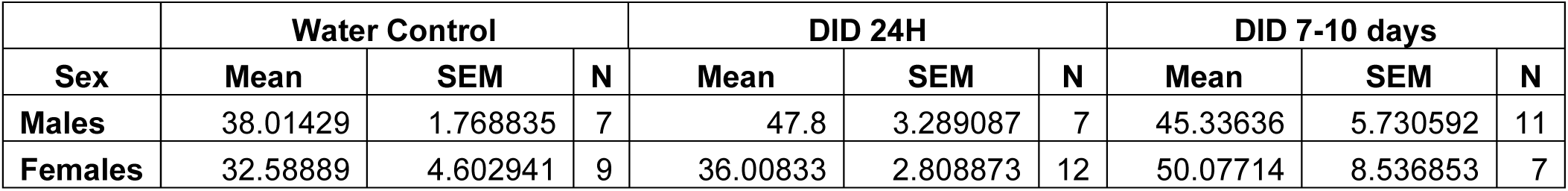
Mouse Membrane Resistance Summary Statistics.

|  | Water Control |  |  | DID 24H |  |  | DID 7-10 days |  |  |
| --- | --- | --- | --- | --- | --- | --- | --- | --- | --- |
| Sex | Mean | SEM | N | Mean | SEM | N | Mean | SEM | N |
| Males | 38.01429 | 1.768835 | 7 | 47.8 | 3.289087 | 7 | 45.33636 | 5.730592 | 11 |
| Females | 32.58889 | 4.602941 | 9 | 36.00833 | 2.808873 | 12 | 50.07714 | 8.536853 | 7 |

**Table 2.** Mouse Membrane Capacitance Summary Statistics.

|  | Water Control |  |  | DID 24H |  |  | DID 7-10 days |  |  |
| --- | --- | --- | --- | --- | --- | --- | --- | --- | --- |
| Sex | Mean | SEM | N | Mean | SEM | N | Mean | SEM | N |
| Males | 144.4943 | 13.39763 | 7 | 140.39 | 7.312513 | 7 | 150.7009 | 11.46159 | 11 |
| Females | 172.0422 | 17.29719 | 9 | 160.6975 | 10.99863 | 12 | 138.8414 | 15.96976 | 7 |

### 5-HT inhibition of OFC neurons is differentially affected in male and female rhesus macaques

Next, we assessed whether the impact of ethanol on 5-HT modulation in the OFC seen in mice generalized to rhesus macaques with a history of ethanol consumption. We recorded whole-cell current-clamp electrophysiological recordings in individual OFC pyramidal neurons from acutely-prepared slices as previously described (Pleil et al., 2024, 2016, 2015) (Fig. 2A, B). Serotonin bath application resulted in a significant interaction of sex and drinking category on change in membrane potential with a main effect of drinking category (HD vs LD vs BD) and the female in the high drinking group having a larger increase in membrane potential in response to 5-HT compared to the low or binge drinking groups. (Fig. 2C). In addition, a regression analysis between 5-HT regulation of membrane potential and life-time consumption of ethanol revealed a significant positive correlation in female, but not male monkey OFC (Fig 2D, E).

**Fig 2.**
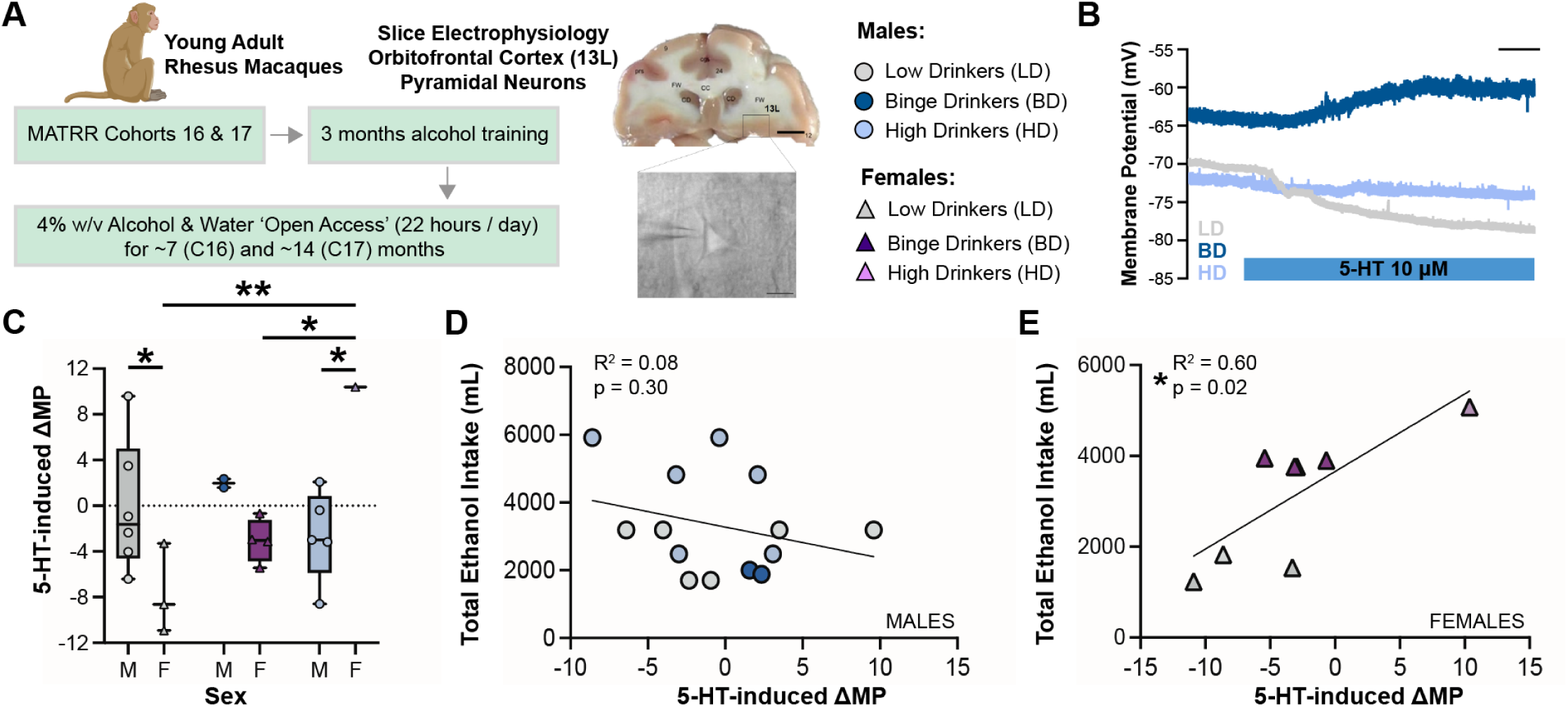
Binge drinking ethanol differentially impacts serotonin signaling in the OFC of rhesus macaques. **A.** Experimental design schematic with tissue block containing area 13L (OFC) in the rhesus macaque and image of a pyramidal neuron recorded from in slice, as viewed under 40X magnification; scale bar: 20 μm. **B.** Representative average membrane potential of low drinking (LD), heavy drinking (HD), and binge drinking (BD) male monkey at 2 min before and 7 min post-application of 5-HT. Scale: 1 min. **C.** Average membrane potential of low drinking (LD), heavy drinking (HD), and binge drinking (BD) male monkey at 2 min before and 7 min post-application of 5-HT (Two-way ANOVA, sex x virus interaction, F_(2,16)_ = 6.396, p = 0.009; virus main effect, F_(2,16)_ = 4.395, p = 0.03; sex main effect, F_(1,16)_ = 0.005, p = 0.945; Sidak post hoc tests, LD males vs females: p = 0.0260, female LD vs HD: p = 0.0072, female BD vs HD: p = 0.0401, HD males vs females: p = 0.0203). **D.** Correlation between total lifetime ethanol intake (in mL) and 5-HT-induced change in membrane potential in all male (Pearson correlation, R^2^ = 0.08, p = 0.30) and **E.** all female monkeys (Pearson correlation, R^2^ = 0.60, p = 0.02). 15 cells (n = 7 LD, n = 6 HD, n = 2 BD) from male monkeys were recorded (n = 2 LD males, 3 HD males, 1 BD males). 8 cells (n = 3 LD, n = 1 HD, n = 4 BD) from female monkeys were recorded (n = 3 LD females, 1 HD females, 3 BD females).

### Ethanol decreases 5-HT_1A_, but not 5-HT_2A_ mRNA levels in the OFC

Our electrophysiological recordings highlighted that there are ethanol-induced changes in serotonin signaling in the OFC. However, it was unclear what serotonin receptor may be responsible for this change. Thus, we used fluorescent *in situ* hybridization (FISH) to visualize and quantify the expression of the 5-HT_1A_ and 5-HT_2A_ receptors in the OFC of water or DID mice (Fig. 3A), as they have previously been shown to modulate OFC function. We found that, compared to water control mice, male and female ethanol mice expressed significantly less *Htr1a* mRNA (Fig. 3B, C). Interestingly, there was a significant positive correlation between *Htr1a* expression and ethanol intake in female mice, but males displayed a trend towards a significant negative correlation (Fig. 3D). Next, we visualized *Htr2a* mRNA and observed no significant difference between water and ethanol mice in either sex (Fig. 3E, F). Finally, we quantified the density of cells that co-expressed *Htr1a* and *Htr2a* mRNA and found significantly less co-expression in ethanol males and females compared to water controls (Fig. 3G, H). We therefore focused our attention on the 5-HT_1A_ receptor.

**Figure 3.**
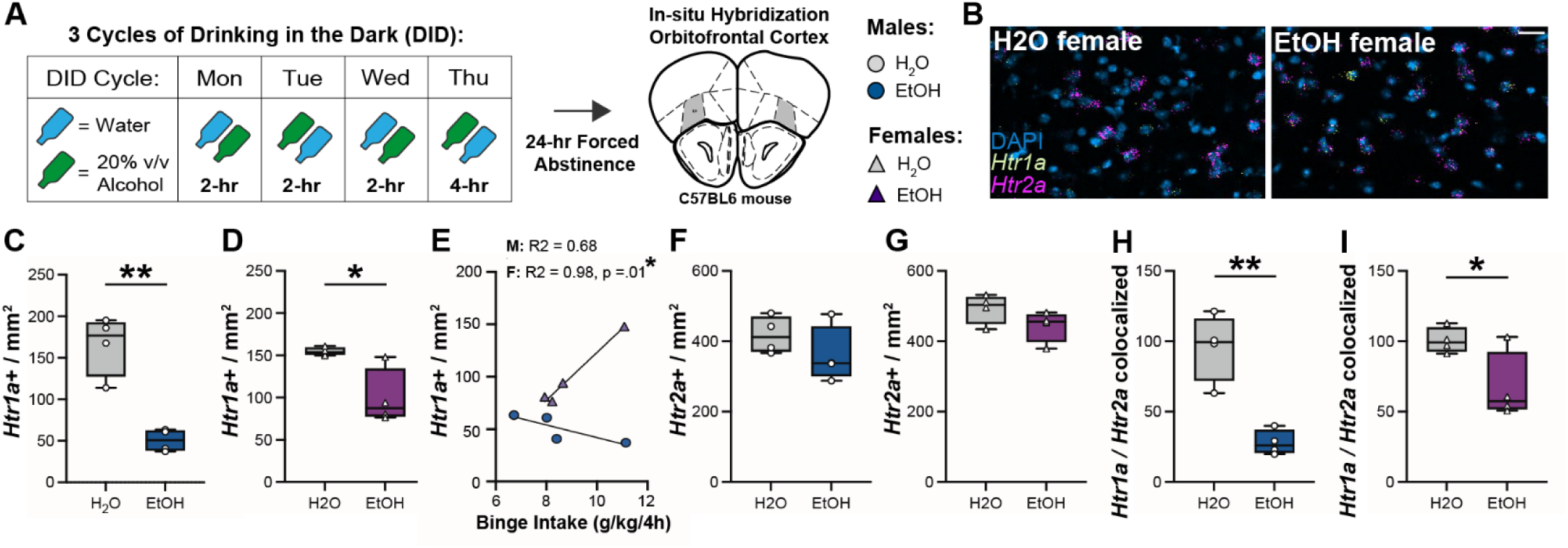
Ethanol drinking reduces expression of the 5-HT_1A_ receptor in the OFC. A. Experimental Timeline. **B.** Representative images of fluorescent *in situ* hybridization to visualize DAPI (blue), *Htr1a* mRNA (yellow), and *Htr2a* mRNA (magenta) in an ethanol-drinking and water control female. Scale: 50 μm. **C, D.** Quantification of *Htr1a* mRNA per mm^2^ of fluorescent image in male and female mice (Unpaired t-test, males: t_(6)_ = 5.931, p = 0.001, females: t_(6)_ = 3.29, p = 0.017). **E.** Correlation between ethanol intake and expression of *Htr1a* mRNA in male and female mice (Pearson correlation, males: R^2^ = 0.684, p = 0.173, females: R^2^ = 0.979, p = 0.01). **F, G.** Quantification of *Htr2a* mRNA per mm^2^ of fluorescent image in male and female mice (Unpaired t-test, males: t_(6)_ = 1.179, p = 0.283, females: t_(6)_ = 1.632, p = 0.154). **H, I.** Quantification of density of cells co-expressing *Htr1a* and *Htr2a* mRNA in the OFC in male and female water or ethanol mice (Unpaired t-tests, males: t_(6)_ = 5.28, p = 0.002, females: t_(6)_ = 2.571, p = 0.042). n = 4 male H2O, n = 4 male EtOH, n = 4 female H2O, n = 4 female EtOH. * p < 0.05; ** p < 0.001

### 5-HT1A-R knockdown in the OFC increases ethanol drinking in males

Our FISH results indicated that 3 cycles of DID led to a reduction in the expression of the 5-HT_1A_ receptor in the OFC. We tested the role of this receptor in behavior using a transgenic mouse to conditionally knock down the 5-HT_1A_ receptor in the OFC. We injected either AAV5-*CamkIIa*-Cre or AAV5-*CamkIIa-*eGFP bilaterally in the OFC of male and female 5-HT1A^fl/fl^ mice (Fig. 4A). Following a recovery period, all mice then underwent a three-week DID protocol and then we tested the impact of OFC 5-HT_1A_ deletion on the consumption of ethanol and sucrose (Fig. 4B). On the third cycle of DID, Cre male mice drank significantly more ethanol compared to control mice during the first two-hour binge session but did not significantly differ in their ethanol preference (Fig. 4C). Throughout the total four hours of the binge drinking session, male Cre mice also drank significantly more ethanol compared to control males (Fig. E, F). Deletion of the 5-HT_1A_ receptor in the OFC did not impact ethanol drinking or preference in female mice. Next, we tested the impact of 5-HT_1A_ receptor deletion on consumption of another rewarding substance, 2% sucrose. Across a 4-h session, female Cre mice drank significantly less sucrose, but did not display a significant difference in preference compared to controls. (Fig. 4G, H).

**Figure 4.**
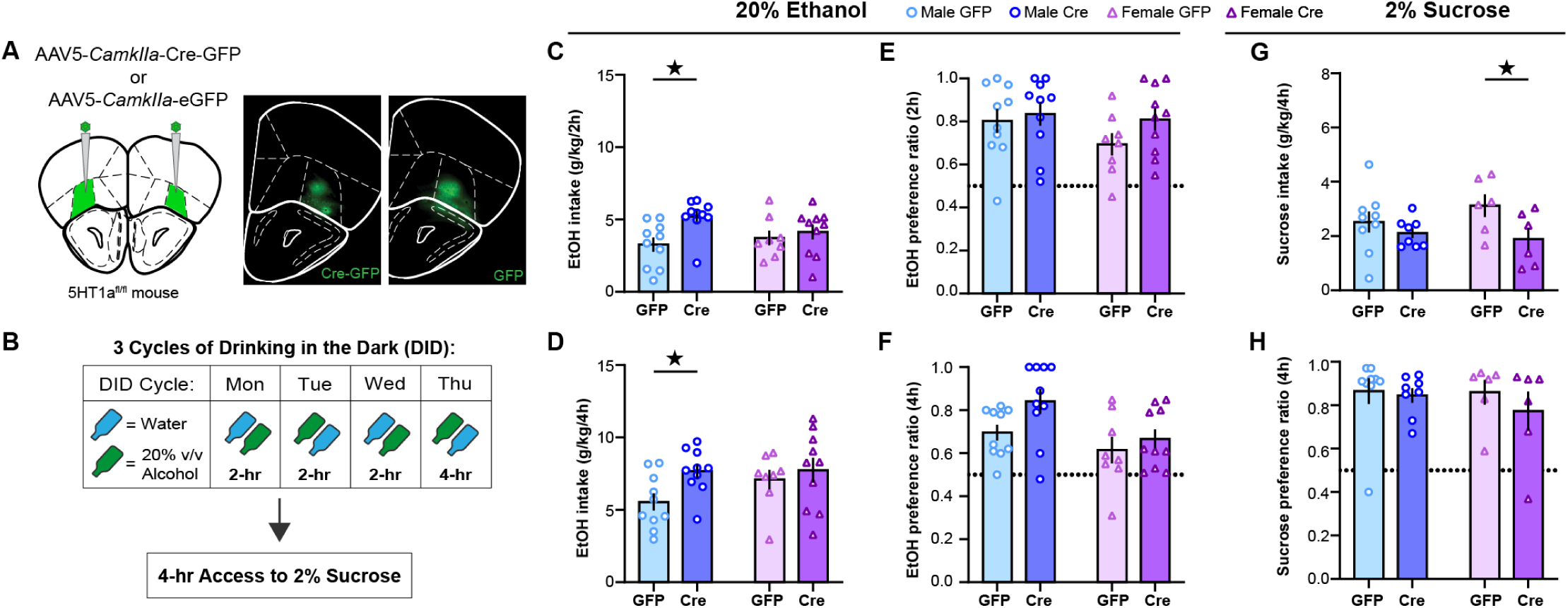
The 5-HT_1A_ receptor in the OFC regulates binge-like ethanol intake in male mice. **A.** Schematic of viral injections of AAV5-*CamkIIa*-Cre-GFP or AAV5-*CamkIIa*-GFP into bilateral OFC of 5-HT1A^fl/fl^ mice and representative images of fluorescent viral expression in the OFC of Cre and GFP control mice. Mice were kept in the home cage for 6 weeks to allow for viral expression. Scale: 500 μm. **B.** Timeline of experiment. **C.** Ethanol intake in g/kg of GFP and Cre male and female mice across the first two hours of the final binge drinking session (Two-way ANOVA, virus main effect, F_(1,34)_ = 6.174, p = 0.018, Sidak’s post hoc, male GFP vs male Cre, p = 0.005; sex main effect, F_(1,34)_ = 0.424, p = 0.519; interaction effect, F_(1,34)_ = 2.645, p = 0.113). **D.** Ethanol intake in g/kg of GFP and Cre male and female mice across the four-hour binge drinking session (Two-way ANOVA, virus main effect, F_(1,34)_ = 4.267, p = 0.047, Sidak’s post hoc, male GFP vs male Cre, p = 0.027; sex main effect, F_(1,34)_ = 0.424, p = 0.519; interaction effect, F_(1,34)_ = 2.645, p = 0.113). **E.** Ethanol preference ratio of GFP and Cre male and female mice across the first two hours of the binge drinking session (Two-way ANOVA, virus main effect, F_(1,34)_ = 1.877, p = 0.18; sex main effect, F_(1,34)_ = 1.54, p = 0.223; interaction effect, F_(1,34)_ = 0.578, p = 0.452). **F.** Ethanol preference ratio of GFP and Cre male and female mice across the four-hour binge drinking session (Two-way ANOVA, virus main effect, F_(1,34)_ = 3.898, p = 0.057; sex main effect, F_(1,34)_ = 6.531, p = 0.015; interaction effect, F_(1,34)_ = 0.869, p = 0.358). **G.** 2% sucrose intake in g/kg of GFP and Cre male and female mice across a four-hour session (Two-way ANOVA, virus main effect, F_(1,25)_ = 5.143, p = 0.032, Sidak’s post hoc, female: GFP vs Cre, p = 0.03; sex main effect, F_(1,25)_ = 0.278, p = 0.603; interaction effect, F_(1,25)_ = 1.335, p = 0.259). **H.** Sucrose preference ratio of GFP and Cre male and female mice across the four-hour session (Two-way ANOVA, virus main effect, F_(1,25)_ = 0.773, p = 0.388; sex main effect, F_(1,25)_ = 0.401, p = 0.532; interaction effect, F_(1,25)_ = 0.294, p = 0.592). * p < 0.05; ** p < 0.001

### 5-HT1A in the OFC is necessary for quinine suppression of ethanol-drinking

As ethanol has been reported to have a bitter taste component, we tested the necessity of the 5-HT_1A_ receptor in the OFC on consumption of quinine water, a bitter aversive tastant. In a new group of mice, we expressed AAV5-*CamkIIa*-Cre or AAV5-*CamkIIa-*eGFP bilaterally in the OFC of male and female 5-HT1A^fl/fl^ mice (Fig. 5A). We then tested mouse intake and preference of 100 μM quinine in water compared to plain water. Deletion of the 5-HT_1A_ receptor from the OFC did not significantly impact intake or preference for quinine across the two- or four-hour session in male or female mice, though female mice did drink more quinine water than males (Fig. 5B-E). Finally, we examined the necessity of the 5-HT_1A_ receptor on consumption of quinine adulterated ethanol as a measure of aversion-resistant drinking. Female Cre mice displayed significantly increased quinine ethanol consumption compared to controls, while male Cre mice displayed an increased preference for quinine ethanol compared to controls. Female mice drank significantly more and had an increased preference for quinine ethanol compared to males. (Fig. 5F, G). Across the four-hour session, male Cre mice drank significantly more quinine adulterated ethanol than control males, while female Cre mice displayed a significantly higher preference for quinine ethanol compared to control females. Female mice drank significantly more and had a higher preference for the quinine ethanol across the four-hour session (Fig 5H, I).

**Figure 5.**
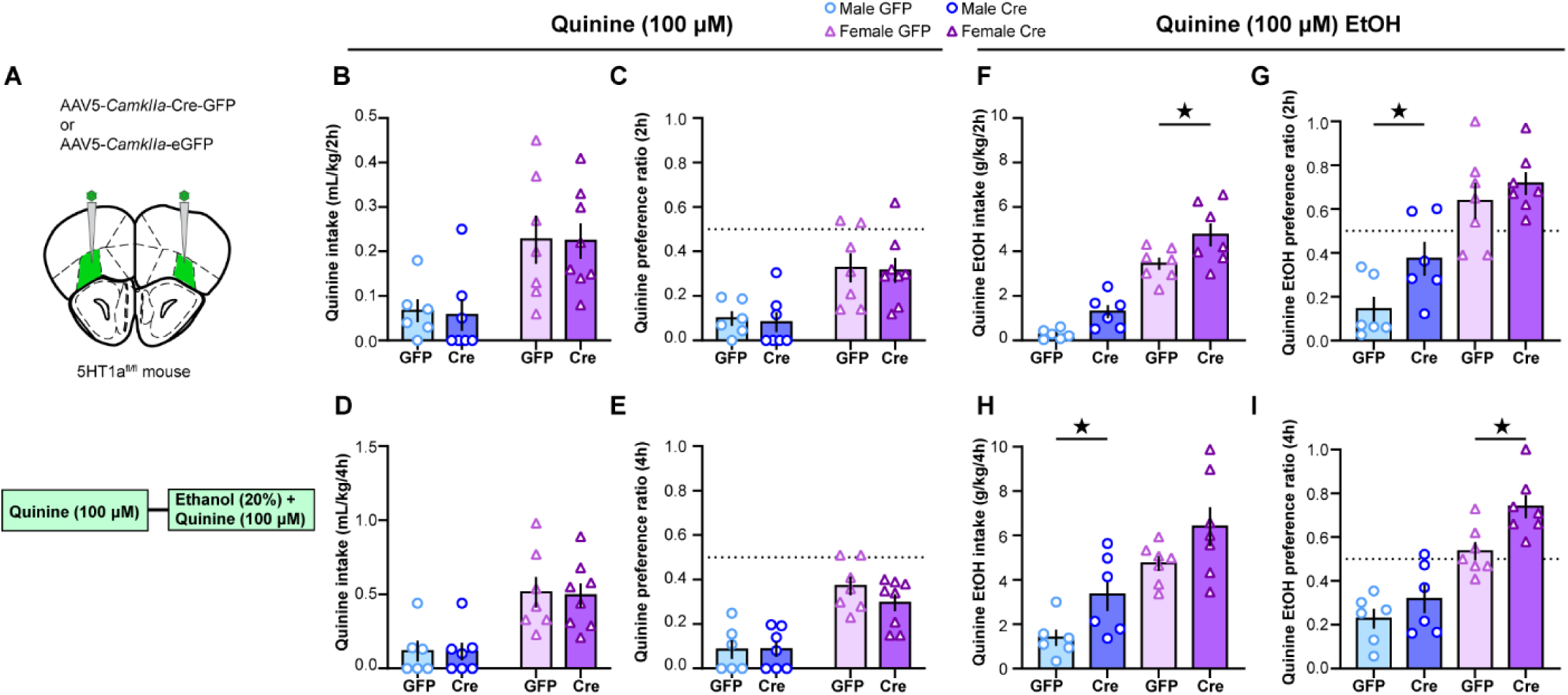
The 5-HT_1A_ receptor in the OFC regulates aversion-resistant ethanol consumption. **A.** Schematic of viral injections of AAV5-*CamkIIa*-Cre-GFP or AAV5-*CamkIIa*-GFP into bilateral OFC of 5-HT1A^fl/fl^ mice and experimental timeline. **B.** Quinine intake in mL/kg of GFP and Cre male and female mice across the first two hours of the four-hour session (Two-way ANOVA, virus main effect, F_(1,24)_ = 0.024, p = 0.877; sex main effect, F_(1,24)_ = 15.636, p < 0.001; interaction effect, F_(1,24)_ = 0.005, p = 0.942). **C.** Quinine preference ratio of GFP and Cre male and female mice across the first two hours of the four-hour session (Two-way ANOVA, virus main effect, F_(1,24)_ = 0.074, p = 0.787; sex main effect, F_(1,24)_ = 18.912, p < 0.001; interaction effect, F_(1,24)_ = 0.002, p = 0.966). **D.** Quinine intake in mL/kg of GFP and Cre male and female mice across the four-hour session (Two-way ANOVA, virus main effect, F_(1,23)_ = 0.001, p = 0.982; sex main effect, F_(1,23)_ = 20.206, p < 0.001; interaction effect, F_(1,23)_ = 0.049, p = 0.827). **E.** Quinine preference ratio of GFP and Cre male and female mice across the four-hour session (Two-way ANOVA, virus main effect, F_(1,23)_ = 1.019, p = 0.323; sex main effect, F_(1,23)_ = 38.179, p < 0.001; interaction effect, F_(1,23)_ = 0.695, p = 0.413). **F.** Quinine ethanol intake in mL/kg of GFP and Cre male and female mice across the first two hours of the four-hour session (Two-way ANOVA, virus main effect, F_(1,22)_ = 11.082, p = 0.003, female GFP vs male Cre, p = 0.012; sex main effect, F_(1,22)_ = 90.105, p < 0.001; interaction effect, F_(1,22)_ = 0.170, p = 0.684). **G.** Quinine ethanol preference ratio of GFP and Cre male and female mice across the first two hours of the four-hour session (Two-way ANOVA, virus main effect, F_(1,22)_ = 5.026, p = 0.035, male GFP vs male Cre, p = 0.034; sex main effect, F_(1,22)_ = 36.917, p < 0.001; interaction effect, F_(1,22)_ = 1.170, p = 0.291). **H.** Quinine ethanol intake in mL/kg of GFP and Cre male and female mice across the four-hour session (Two-way ANOVA, virus main effect, F_(1,22)_ = 8.031, p = 0.010, male GFP vs male Cre, p = 0.049; sex main effect, F_(1,22)_ = 25.32, p < 0.001; interaction effect, F_(1,22)_ = 0.051, p = 0.823). **I.** Quinine ethanol preference ratio of GFP and Cre male and female mice across the four-hour session (Two-way ANOVA, virus main effect, F_(1,22)_ = 8.215, p = 0.009, female GFP vs Cre, p = 0.008; sex main effect, F_(1,22)_ = 50.692, p < 0.001; interaction effect, F_(1,22)_ = 1.208, p = 0.283). Quinine only: n = 6 GFP males, n = 7 Cre males, n = 7 GFP females, n = 8 Cre females; Quinine ethanol: n = 6 GFP males, n = 6 Cre males, n = 7 GFP females, n = 7 Cre females.

## DISCUSSION

The orbitofrontal cortex (OFC) is important for decision making and controlling goal-directed behavior and is disrupted in AUD and substance use disorders (SUD). Here, we wanted to test the impact of binge-like ethanol consumption on 5-HT modulation of OFC physiology and then determine how this contributes to behavior. 5-HT hyperpolarizes the OFC in naïve mice, but chronic intermittent ethanol vapor exposure leads to a blunting of this hyperpolarizing effect during forced abstinence (Nimitvilai et al., 2018, 2017, 2016). Thus, we first explored whether voluntary binge drinking also alters 5-HT modulation of the OFC. First, we recorded from neurons from mice. While 5-HT hyperpolarizes OFC neurons from control male mice, binge-like ethanol consumption reduced this hyperpolarization. Interestingly, neurons from female water mice were not hyperpolarized in response to 5-HT and there were no significant differences in the change of resting membrane potential in female mice that drank ethanol. This suggests a fundamental difference in how 5-HT can modulate function in the OFC, consistent with previous reports from our group where we found sex differences in the regulation of 5-HT function in the bed nucleus of the stria terminalis and the lateral habenula (Flanigan et al., 2023b). However, this does stand in contrast to a recent report suggesting no differences in 5-HT modulation of OFC function in rats (Mooney et al., 2025). The DID protocol requires mice to be individually housed for accurate alcohol consumption measurements, however, female rodents are more susceptible to social isolation stress which could be a contributing factor to the female controls showing a lack of hyperpolarization after 5-HT. There are additional differences, including animal species, that may drive this lack of generality.

We next shifted to recording from acute brain slices obtained from rhesus macaques to translate our findings across species that may more closely model the human brain. While we expected to observe a similar effect as mice of ethanol drinking on 5-HT’s effects on resting membrane potential, the effect in monkeys was different. While water-drinking male mice displayed 5-HT-induced hyperpolarization, male low ethanol drinking monkeys did not show a change in membrane potential after application of 5-HT. This suggests that even low levels of alcohol consumption over long periods of time may alter 5-HT regulation of the OFC, however, no true water controls are used in these MATRR cohorts, limiting the generality of these findings. In contrast neurons from female monkeys displayed 5-HT-induced hyperpolarization. Interestingly, the monkeys are allowed periods of socialization, which may be protective for female monkeys. Male mice that were 24 hours or 7-10 days post last alcohol drinking session had an inhibition of hyperpolarization after 5-HT application, which was significantly different than the water drinking males. Female ethanol-drinking mice did not display a significant difference from water-drinking mice. While sample sizes for the monkey recordings are small, interesting differences between groups were observed. Notably, we found that there was a main effect of drinking status on 5-HT-induced changes in membrane potential. However, because of the nature of these experiments, the data was not distributed evenly across drinking categories. We next directly assessed whether the total amount of lifetime EtOH consumed correlated with 5-HT-induced changes in membrane potential. Interestingly, we found the greater their ethanol intake the more likely 5-HT was to increase the membrane potential. One interpretation is that in monkeys with greater 5-HT_1A_-R effects there is reduced ethanol consumption. It also could be related to 5-HT_2A_-R, as monkeys with greater 5-HT2A-R driven depolarization may consume more ethanol. It is important to consider that there was no water drinking control group in the monkeys, which limits comparison between mice and monkeys.

Our electrophysiological recordings indicated a shift in neuronal response to serotonin after binge-like ethanol consumption. Thus, we wanted to examine which serotonin receptor may be responsible for this change. We used FISH and observed a significant reduction in expression of the 5-HT_1A_, but not the 5-HT_2A_ receptor, in the OFC of male and female mice. Our initial hypothesis was that we would observe a decrease in 5-HT_1A_ receptor with a concomitant increase in expression of the 5-HT_2A_ receptor. Previous studies using chronic intermittent ethanol found increases in excitability of OFC neurons and reduced 5-HT-induced inhibition of OFC neurons via an impact on the 5-HT_1A_ receptor (Nimitvilai et al., 2018, 2017, 2016). Further, it has been reported that cocaine exposure can downregulate 5-HT_1A_ receptor and upregulate 5-HT_2A_ receptor expression in the OFC (Wright et al., 2017). Therefore, the lack of change in the 5-HT_2A_ receptor was surprising but revealed a potentially important drug-specific change in the 5-HT system.

The reduction in 5-HT_1A_ receptor mRNA we observed in binge-like ethanol drinking led us to test the impact of 5-HT_1A_ receptor deletion from the OFC. During a binge-like ethanol drinking session, male mice with deletion of 5-HT_1A_ receptor (Cre males) had increased ethanol intake and preference compared to control males. This suggests that 5-HT_1A_ receptor signaling in the OFC may constrain ethanol consumption under normal conditions. This is especially interesting in light of a recent study demonstrating that there is an ensemble of OFC neurons that is important for inhibiting ethanol consumption (Gimenez-Gomez et al., 2025). It raises the intriguing idea that perhaps 5-HT function is important for shaping the engagement of this ensemble. In comparison to the male mice, female mice with deletion of 5-HT1A showed no significant difference from control females in ethanol intake or preference. This presents the possibility that 5-HT signaling in the OFC could contribute to binge alcohol drinking in males, but not females, or perhaps a different 5-HT receptor is engaged. However, this does highlight the importance of considering sex in development of new pharmacological tools.

Ethanol is a rewarding substance that mice voluntarily drink and that acts on the mesolimbic reward system in the brain. Because we found that deletion of the 5-HT_1A_ receptor impacted ethanol drinking, we next wanted to test whether this effect was generalizable to non-ethanol rewards. Interestingly, deletion of 5-HT_1A_ decreased sucrose consumption, but only in female mice. This suggests that 5-HT_1A_ in the OFC may be important for regulation of non-ethanol rewards in females and is specific to regulation of ethanol reward in males. Future work should examine if the 5-HT_1A_ receptor is essential for intake of other non-ethanol drugs. While the OFC and 5-HT are implicated in reward, they are also implicated in aversive processing and the OFC in particular is important for reward valuation (Rolls et al., 2020). We tested the necessity of the 5-HT_1A_ receptor in consumption of an aversive substance, quinine, which has a bitter taste. Deletion of OFC 5-HT_1A_ did not significantly impact quinine intake or preference in male or female miceThis suggests that this receptor is not mediating aversion alone. Ethanol is a reward, but alcohol abuse/ AUD leads to many negative consequences as well—social problems, hangover, physical dependence, financial issues, and work performance. Therefore, we tested the impact of 5-HT_1A_ deletion on aversion-resistant ethanol drinking, by testing intake and preference for quinine adulterated ethanol, a model of consumption despite negative outcomes. Deletion of the 5-HT_1A_ receptor in the OFC increased quinine-ethanol intake and preference in both male and female mice. This suggests that proper function of the 5-HT_1A_ receptor in the OFC is necessary for suppression of aversion-resistant ethanol drinking, which may be related to the OFC’s role in reward valuation. Unlike ethanol alone, this manipulation impacted drinking in male and female mice, suggesting that the 5-HT_1A_ receptor in the OFC may be particularly important in AUD where binge drinking is leading to negative consequences and may have a difference in reward value.

Overall, we have identified a mechanism by which binge-like ethanol consumption impacts 5-HT signaling in the OFC. Binge-like ethanol drinking reduces the expression of the 5-HT_1A_ receptor, which normally hyperpolarizes neurons upon activation via 5-HT. This loss of 5-HT_1A_ receptor prevents inhibition of binge-like ethanol drinking and aversion-resistant ethanol drinking. The 5-HT_1A_ receptor represents a promising target for therapeutic interventions for AUD, as increasing expression or efficiency of this receptor may prevent excessive ethanol intake.

## AUTHOR CONTRIBUTIONS

M.P., T.K., K.A.G. conceptualized the experiments. M.P., S.D., and K.M.B. conducted the experiments. M.P., J.A.W., J.L.R. analyzed data. M.P. and J.A.W. created figures. J.A.W., M.P. and T.K. wrote the paper with editing contributions from all authors.

## ACKNOWLEDGMENTS

This work was supported by National Institutes of Health K12 GM000678 (to J.A.W.), AA028298, AA026485 (to M.M.P.), AA019454 (T.L.K.) and AA01943, OD011092-66 (K.A.G.) We thank the Division of Comparative Medicine (DCM) at the University of North Carolina at Chapel Hill for assistance with rodent husbandry, breeding, and veterinary support.

